# Accelerating Natural Product Discovery with Linked MS-Genomics and Language/Transformer-Based Models

**DOI:** 10.1101/2025.01.20.633932

**Authors:** Dillon W. P. Tay, Winston Koh, Shi Jun Ang, Zicong Marvin Wong, Yi Wee Lim, Elena Heng, Naythan Z. X. Yeo, Krishnan Adaikkappan, Fong Tian Wong, Yee Hwee Lim

## Abstract

An integrated multi-modal characterization of a microbial strain library streamlines the effort for natural product discovery. By integrating language- and transformer-based models to cross-validate mass spectrometry (MS)-genome datasets, microbial producers of diverse natural products are rapidly identified with high (75-100%) precision. Our findings demonstrate the transformative potential of linked MS-genome datasets at the strain-level to significantly accelerate discovery and enhance our understanding of microbes beyond currently known and curated knowledge.

## Main

Natural products (NPs) offer exceptional chemical diversity and therapeutic potential^1^. However, their discovery is hindered by inefficiencies in linking genomic information to structural validation^2^. Despite advancements in genomic and analytical technologies such as mass spectrometry (MS), existing approaches often rely heavily on predefined rules and reference libraries^3^, limiting comprehensive exploration of natural chemical space.

To address these challenges, we introduce a framework utilizing integrated language- and transformer-based models to cross-validate and rank linked MS-genome datasets to enable scalable inference of secondary metabolites from microbial strain libraries. Protein language models (ESM2)^4^ efficiently encode genomic data into meaningful embeddings, overcoming limitations of fragmented genomic sequences, while an in-house developed workflow known as WISE (Workflow for Intelligent Structural Elucidation), leverages molecular language processing and transformer-based structural elucidation models for direct interpretation of MS/MS data to structures. This integrative approach significantly accelerates the identification of microbial producers and corresponding compounds, bridging the gap between data generation and actionable insights with increased confidence, thereby streamlining the natural product discovery pipeline (**Figure 1A**).

**Figure 1.**
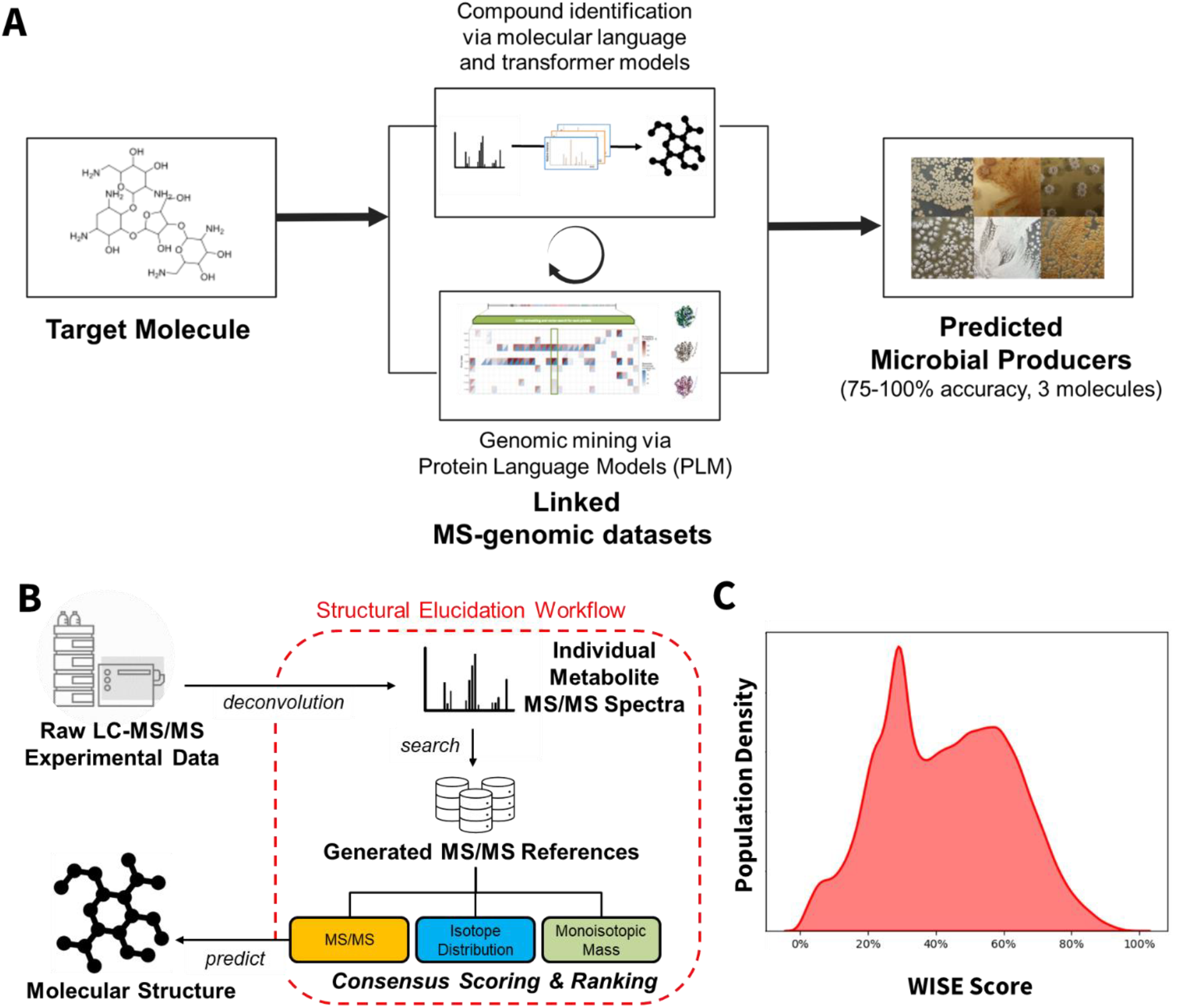
**(A)** Concept of a linked genomics and mass spectrometry (LCMS/MS) multi-modal dataset and predictive models for streamlined discovery of natural products in microbial strain libraries. Putative producers of compounds of interest are identified through co-occurrence of the best ranked strains via protein language models embeddings and the highly ranked strains predicted to produce the target molecules via our in-house developed tandem mass spectral MS/MS to structural elucidation workflow (WISE). **(B)** Overview of the cheminformatics analysis workflow (WISE) to annotate individual molecules from raw liquid chromatography-tandem mass spectral (LC-MS/MS) data of fermentation extracts. **(C)** Distribution of confidence scores (WISE scores) for 29,112 compounds identified by WISE from the 2,138 curated LC-MS/MS dataset.

Natural product discovery traditionally suffers from bottlenecks associated with limited reference MS/MS databases^5^, restricting identification mostly to characterized metabolites^6^. To address this critical bottleneck, we describe the integration of three technologies comprising (1) molecular language processing, (2) structure-to-MS/MS models, and (3) a data processing workflow, synergistically designed to directly elucidate molecular structures from LC-MS/MS spectra of mixtures (e.g., fermentation extracts). To enable the identification of novel structures, we leverage molecular language processing, a generative technology that simulates the “language” of molecular structures, where more than 67 million known and novel natural products-like compounds were previously sampled from the trained model^7^. Predictive transformer-based structure-to-MS/MS models^8^ for reference spectra generation were also fine-tuned to include collision energy as a parameter. By separately generating spectra within ±5 eV collision energies from each other, we improved cosine similarity between predicted and experimental MS/MS spectra from an average of 59% to 75%. With an average alignment of 75% with experimental spectra, we have improved reference reliability and accuracy (42%) in computational structural elucidation significantly compared to state-of-the-art methods^8-10^ (8∼13% accuracy) (see **Table S1** for comparison).

By predicting spectral references for novel molecules generated through molecular language processing, we address limitations posed by sparse spectral references in current databases, offering a robust and rapid method for identifying novel compounds in an expanded chemical exploration space. To further translate these capabilities into actionable insights, we developed a data processing workflow named WISE (Workflow for Intelligent Structural Elucidation) that leverages the generated reference MS/MS database to directly interpret complex LC-MS/MS data of mixtures into its constituting molecular structures in a scalable discovery process (**Figure 1B**). In this study, WISE is deployed to deconvolute (**Figure S1**) and annotate the 2,138 distinct LC-MS/MS dataset^11^ generated from a diverse set of 54 actinobacteria and their mutants from the A*STAR National Organism Library^12^ using an integrase-mediated multi-faceted approach^13^. Using this method, we identified 29,112 unique compounds with an average WISE score of 43% (**Figure 1C**, see methods for WISE score definition). Notably, different processing algorithms have enabled WISE to significantly enhance the scale of identification, resulting in the number of unique compounds identified through WISE being nearly 40 times greater than those previously processed using GNPS molecular networking analysis. While the GNPS workflow^6^ groups spectra based on their MS/MS similarity, WISE conducts structural identification for individual spectra, which has the potential to yield a higher number of unique annotations.

In parallel, recent advances in protein language models (PLMs) have significantly enhanced *in silico* biophysical analysis, especially in structure-related tasks such as secondary structure prediction^4^ and contact prediction^14^. Embeddings from these models can extract remote homology information that may be obscured in primary sequences, enabling a more sophisticated understanding of protein function^15,16^. Unlike sequence homology, which emphasises conserved regions, these embeddings represent proteins as high-dimensional vectors, capturing structural and functional nuances that can extend beyond sequence similarity^15,16^. These high-resolution embeddings can subsequently facilitate rapid similarity searches through vector databases such as Faiss^17^, allowing high-confidence, scalable, and efficient genomic analyses. Building on these advancements, we chose to concentrate on higher-resolution protein-level embeddings instead of those derived from entire biosynthetic gene clusters (BGCs). Moreover, this approach circumvents the necessity for high-quality genome assemblies, as protein sequences can be annotated even from fragmented or incomplete genomes, as demonstrated in our current genomic dataset (**Table S2**).

To address the challenges in overfitting of these individual models, we implemented a co-occurrence prioritisation strategy that combines and validates protein language model embeddings from genomic data with WISE scores derived from MS/MS signals. When applied to our strain library^13^, this multi-modal cross-validation strategy facilitates rapid and highly confident identification of microbial producers of diverse natural products – valinomycin (**Figure S2, Table S6**), surfactin B (**Figure S3, Table S7**) and neomycin B (**Figure 2, Table S8**).

**Figure 2.**
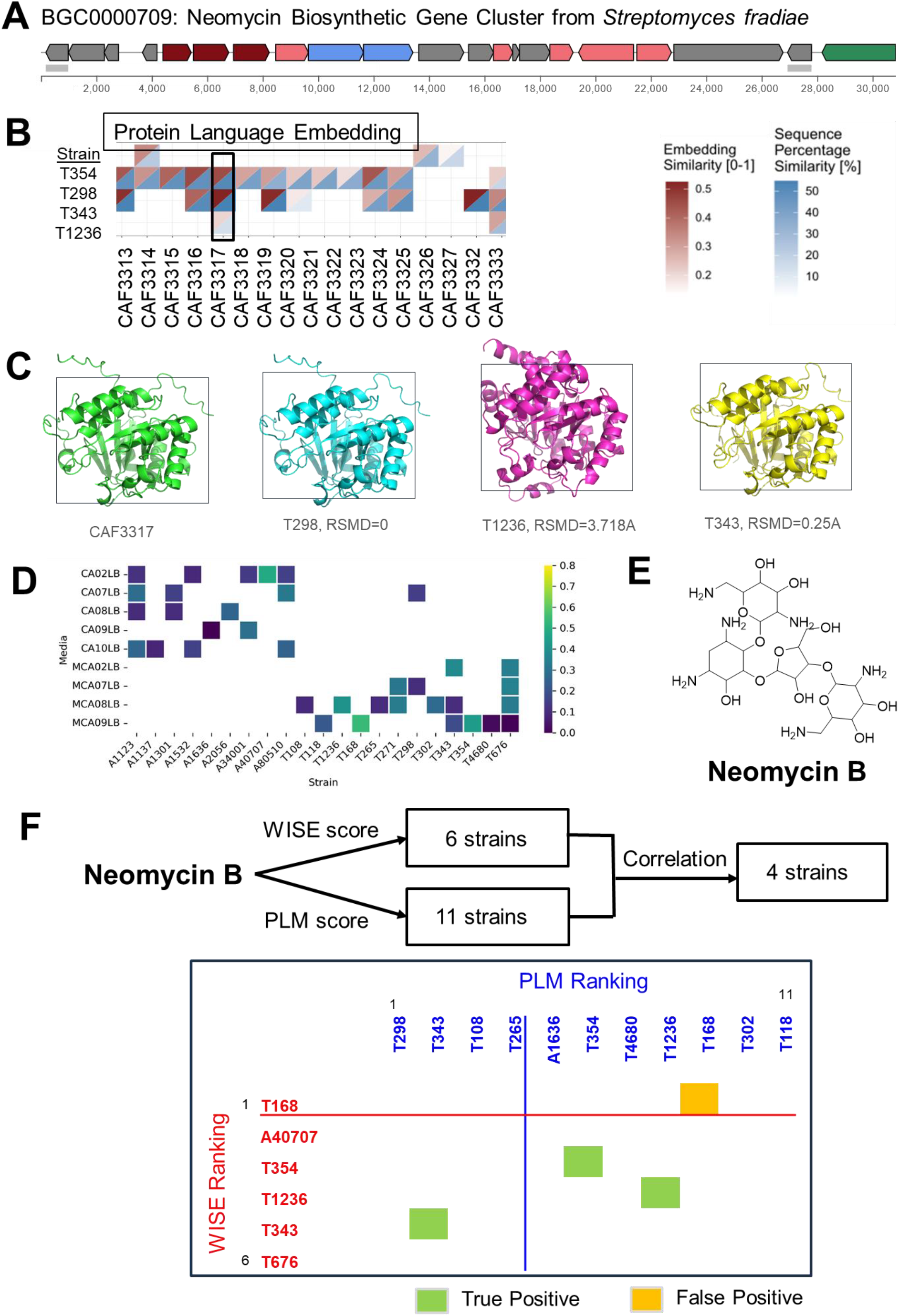
Case Study: Neomycin B. **(A)** Elucidated biosynthetic gene cluster BGC00007089, responsible for neomycin production. **(B)** Similarity (%) of each embedded protein and sequence similarity (%) of the top-scoring protein matches. **(C)** Predicted protein structures of the top matches for Fe-S oxidoreductase (CAF33317) from neomycin BGC. **(D)** Heatmap of WISE scores for strain-media combinations derived from the liquid chromatography-tandem mass spectrometry (LC-MS/MS) data of fermentation extracts. Only strains with at least 1 media combination with non-zero WISE score are shown. Activated strains (if any) are referred to by their parent strain instead of individually (details of the activated strains are in **Table S8**). **(E)** Structure of neomycin B. **(F)** Workflow of the co-occurrence strategy and matrix ranking table with 35% WISE score and 0.75 PLM score cutoff. Strains that have been cross validated are highlighted in green or yellow, indicating whether these compounds were present (green) or absent (yellow). Lines indicating higher confidence thresholds of 50% WISE score (in red) and 2.5 PLM score (in blue) are also included.

In the analysis of neomycin B (**Figure 2E**), the BGC encoding neomycin biosynthesis (BGC00007089^18^, **Figure 2A**) was examined using the ESM2 protein language model to embed protein components. These embeddings were compared against a vector database derived from the genomic dataset using nearest neighbour algorithms to identify top matches (**Figure 2B**). Pairwise sequence alignments and structural alignments of predicted structures of the enzyme within the BGC (Fe-S oxidoreductase, CAF33317) also validated the model’s effectiveness for capturing sequence and structural relationships (**Figure 2C**). Separately, the LC-MS/MS data of the strains’ fermentation extracts were also processed via WISE workflow (**Figure 2D**). Highly ranked strains by both cheminformatics (WISE) and bioinformatics (PLM), with a minimum of 35% WISE score or 0.75 PLM score were identified through this cross-validation matrix (**Figure 2F**). 4 unique strains were predicted to be neomycin B producers based on the co-occurrence of PLM and WISE scores, of which three were validated (75% accuracy rate) through authentic product MS/MS spectral matching. Notably, two of the validated species were previously uncharacterized *Streptomyces* producers of neomycin (**Table S5**).

By employing such a ranking score matrix composed of PLM embeddings and WISE scores with cutoffs, producers of two other target compounds (valinomycin and surfactin B, **Figures S2 and S3**) were also identified with accuracies ranging between 86-100%. It is especially noteworthy that the case study results were achieved without any pre-validation against external models, experimental reference MS/MS data, or high-quality genomic information and annotation, demonstrating this integrative approach as a powerful strategy to uncover microbial producers.

Collectively, the multi-modal integration framework significantly addresses several longstanding challenges in natural product research, such as untargeted metabolite inference and identification of new producers in a scalable and accelerated fashion. Individually, advanced techniques in molecular language processing, transformer-based structural predictions, and protein-level embeddings can rapidly identify patterns in their respective datasets. We are cognizant that challenges such as data quality and model interpretability can limit this approach. Predictions based solely on PLM or WISE scores can significantly overestimate potential producers. Therefore, it is critical to ensure biological relevance. Utilizing the co-occurrence strategy with defined confidence thresholds helps achieve high accuracy (75-100%). Through integrating both chemical and genomic insights, we effectively bridge the critical gap between data generation and actionable biological insights, substantially enhancing confidence in producer strains and compound identification. Like with all prediction methodologies, there remain false positives (2/14 for valinomycin and 1/4 for neomycin B) despite consensus in PLM and WISE score alignment, possibly due to data quality and inherent bias in the processing algorithms. The complexity of interpreting high-dimensional data and the potential for error in vector-based searches underscore the need for continued refinement of our algorithms and methodologies. Ultimately, ensuring high quality MS/MS data and robust biosynthetic hypotheses is essential to fully leverage this innovative discovery framework and reduce the rate of missed producers. Nonetheless, our methodology can facilitate rapid discovery of natural products and previously unrecognized producer strains, expanding the boundaries of known chemical and microbial biological diversity.

## ONLINE METHODS

### Genomics sequencing

Genomic libraries were prepared for each strain using 10 ng of genomic DNA and the plexWell™ 96 kit, according to the manufacturer’s instructions. Sequencing was performed on an Illumina HiSeq 4000 platform using a 2 × 151 bp paired-end protocol. Raw Illumina sequencing reads were first evaluated using FastQC to assess base quality, adapter contamination, and sequence duplication. Adapter trimming and quality filtering were performed using Trimmomatic^19^ with a minimum read length of 36 bp. High-quality reads were then assembled de novo using SPAdes with parameters optimized for microbial genome reconstruction. Assembly quality was quantified using N50 and total assembly size, across the dataset. Protein-coding sequences were identified from assembled contigs using Prodigal^20^. The predicted protein sequences within these regions were then extracted to serve as candidates for downstream computational analysis, including embedding-based similarity search.

### Similarity searches Using kNN in protein language model embeddings

To identify functional analogs across the sequenced strains, we applied ESM2, a transformer-based protein language model, to generate embeddings for the predicted proteins. Embeddings were computed following the ESM2 protocol (https://github.com/facebookresearch/esm) and indexed using an OpenSearch vector database, paired with metadata including strain identifiers and environmental origin. Similarity searches were conducted using a k-nearest neighbors (kNN) algorithm based on cosine distance in the high-dimensional embedding space. For each query protein— such as enzymes linked to specific compound classes in the MIBiG reference database— we retrieved the top 20 most similar proteins, ranked by embedding proximity. This approach enabled rapid prioritization of homologous or convergently evolved enzymes potentially involved in natural product biosynthesis, even in the absence of high sequence identity, demonstrating the utility of language model-based representations in functional protein mining. Using a PLM approach on fragmented genomes, the removal of cluster localisation constraints may lead to the possibility that genes are coincidentally found within the same genome but do not work in concert with one another. To overcome this limitation, strains are also ranked in terms of total similar proteins observed from BGC as a means of prioritising possible producers, rather than looking at individual protein hits.

### Structure-to-MS/MS

Tandem mass spectra (MS/MS) of molecular structures were predicted using a transformer-based model adapted from ICEBERG^8^ that takes as inputs: (1) molecular structure in SMILES format, (2) adduct type, and (3) collision energy in eV, to predict their corresponding MS/MS spectral reference. Spectral references were generated for structures from both the COCONUT^21^ and generated 67 million natural product^7^ databases for 4 adduct types – [M+H]^+^, [M+Na]^+^, [M+K]^+^, [M+NH_4_]^+^ and a collision energy range from 1 – 129 eV to create an augmented MS/MS spectral reference database for structural elucidation.

### Workflow for Intelligent Structural Elucidation (WISE)

The workflow begins by performing analyses of MS1 signals at the individual m/z level that separates MS1 signals into three categories: (1) “Noise-like”: signals with similar intensities across the entire time range, indicating a noise or background signal, (2) “Fragment-like”: signals with more than 1 major peak, usually localized around the same retention time, indicating a common scaffold or fragment from compound analogs eluting at different distinct times, and (3) “Compound-like”: signals which follow a regular peak shape distribution, in line with compound elution from the column (**Figure S1**). The MS/MS spectra of the deconvoluted compound-like signals are compared against the generated MS/MS reference database across three metrics of (1) MS/MS spectra similarity, (2) monoisotopic mass difference, and (3) isotopic distribution similarity, to give a WISE score. For each of the deconvoluted compound-like signals, the corresponding precursor *m/z* is first used to shortlist and filter the augmented MS/MS spectral reference database for entries that are within ±0.005 *m/z*. WISE scores are then calculated between the identified compound and every shortlisted reference according to the following formula:

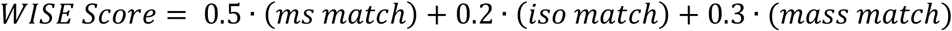

Where,

*ms match* = weighted cosine similarity between experimental MS/MS and generated MS/MS reference

*iso match* = moderated cosine similarity between experimental MS1 isotopic distribution and calculated isotopic distribution

*mass match* = similarity scoring between experimental precursor *m/z* and calculated theoretical monoisotopic mass

Formulas for the individual match scores as follows:

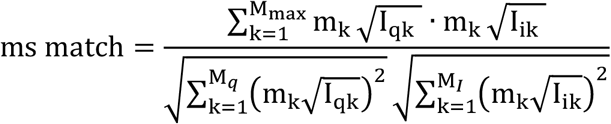

I_q_ and I_i_ are vectors of *m/z* intensities representing the experimental spectrum and the predicted spectrum respectively. m_k_ and I_k_ are the *m/z* ratio and intensity found at *m/z* = k; M_q_ and M_i_ are the largest indices of I_q_ and I_i_ with non-zero values. M_max_ is the larger of M_q_ and M_i_. The cosine similarity is weighted by *m/z* as the peaks in mass spectra corresponding to larger fragments are more characteristic and useful in practice for identifying^22^.

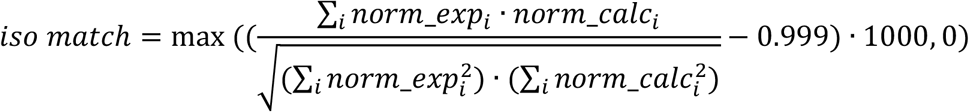

Norm_exp_i_ and norm_calc_i_ are MS1 intensities of the isotopic distributions divided by the maximum MS1 intensity in their distribution of the experimental and calculated isotopic distributions respectively. As the cosine similarity score of the isotopic distribution is inherently high due to the identical *m/z* values (M, M+1, M+2, M+3…etc) and minute differences between low natural abundance of heavier isotopes, the score is moderated to only consider similarities beyond 99.9% (i.e. a cosine similarity of 99.9882% is moderated to 88.2% instead). Anything below the 99.9% threshold is taken as 0%.

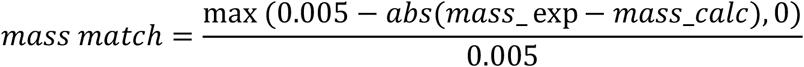

Mass_exp is the precursor *m/z* of the experimental compound observed and mass_calc is the calculated theoretical precursor *m/z* of the reference compound. The previous mass filter of ±0.005 *m/z* is taken here as the maximum possible deviation from the experimental precursor *m/z* (i.e. 0.005 *m/z* difference is 0% mass match).

Note: WISE’s performance heavily depends on data quality. Low abundance metabolites or noisy signals can hinder accurate identification. For instance, low abundance metabolites may produce fewer MS/MS signals or have their signals obscured by random noise, leading to poorer matches with the generated database. This may explain the spike around ∼30% score observed in **Figure 1C**.

### Metabolomic characterization of the strain library

The reported 2,138 LC-MS/MS dataset^13^ is re-processed using WISE that combines data processing, deconvolution, and structural elucidation in an end-to-end automated processing pipeline to detect metabolites present based on their MS1 signal intensities exceeding a threshold of 10,000 in their base peak chromatogram (BPC). The top 10 predicted candidate molecular structures per metabolite identified were taken to form a pool of metabolites from which searches were done to identify producers of valinomycin, surfactin B, and neomycin B.

### Case study validation

The presence of valinomycin and surfactin B in microbial strain fermentation extracts were validated via isolation and structural confirmation by NMR^13^, as well as independent spectral matching of experimental LC-MS/MS data against the Global Natural Products Social Molecular Networking (GNPS)^6^ spectral reference libraries^11^. The library spectra were filtered by removing all MS/MS fragment ions within +/- 17 Da of the precursor *m/z*. MS/MS spectra were window filtered by choosing only the top 6 fragment ions in the +/- 50Da window throughout the spectrum. The precursor ion mass tolerance was set to 0.02 Da and a MS/MS fragment ion tolerance of 0.02 Da. All matches kept between queried experimental spectra and reference library spectra were required to have a cosine similarity score above 0.7 and at least 6 matched peaks.

The presence of Neomycin B in microbial strain fermentation extracts was validated by spectral matching between experimental LC-MS/MS data against an in-house reference MS/MS obtained from analyzing a commercial standard of Neomycin B, as well as independent comparison with the GNPS reference of Neomycin B (CCMSLIB00010114511). Specifically, the WISE processed experimental LC-MS/MS dataset was filtered to those annotated as [M+Na]^+^ adducts, had precursor *m/z* that was ±0.005 *m/z* of 637.3015062321 (the calculated theoretical monoisotopic mass of the [M+Na]^+^ adduct of Neomycin B), and a retention time between 6.94 – 7.14 min (the range of experimental retention times observed for Neomycin B). The shortlist was further filtered to keep only experimental spectra with at least 4 matched peaks (that were at least 5% of base peak abundance and ±5 Da away from each other) to both in-house commercial and GNPS reference spectra.

## DATA AVAILABILITY

The LC-MS/MS dataset of fermentation extracts from 54 actinobacterial strains and their mutants used in this work has been deposited and is publicly accessible via MassIVE with the accession number MSV000092237 (https://doi.org/doi:10.25345/C53X83W53). Reference MS/MS spectra of Valinomycin (CCMSLIB00005721598), Surfactin B (CCMSLIB00005727861), and Neomycin B (CCMSLIB00010114511) are available from the Global Natural Products Social Molecular Networking (GNPS)^6^ spectral reference libraries.

## CODE AVAILABILITY

The code used to train the molecular language processing model as well as the trained model used to generate natural product-like molecules is available from Github at https://github.com/SIBERanalytics/Natural-Product-Generator.^7^ The code used to train the structure-to-MS/MS model (ICEBERG)^8^ as well as the pre-trained model is available from Github at https://github.com/samgoldman97/ms-pred. The data processing Workflow for Intelligent Structural Elucidation (WISE)^23^ is partially based on Mestrenova v15.0.1-35756 which is commercially available from Mestrelab Research S. L. (www.mestrelab.com). The remaining code in WISE for MS/MS-to-structure identification and to recreate the application case studies shown here is available from Github at https://github.com/SIBERanalytics/MS2-to-Structure. The code used for generating protein sequence embeddings using ESM-2, indexing them in a vector database for fast similarity searches via approximate nearest neighbor (ANN) methods is available from Github at https://github.com/lianchye/pLM_analysis.git.

## ACKNOWLEGEMENTS

The authors gratefully acknowledge financial support from the National Research Foundation, Singapore (NRF-CRP19-2017-05-00). This work is also supported by the Agency for Science, Technology and Research (A*STAR, Singapore) under C211917003, C211917006, C233017006, and Singapore Integrative Biosystems and Engineering Research Strategic Research & Translational Thrust (SIBER SRTT). We gratefully acknowledge the Biocomputing Centre at the A*STAR Bioinformatics Institute for their crucial role in hosting and managing the genomics datasets that were fundamental to this research. We also thank the compute support team at the A*STAR Computational Resource Centre (A*CRC) for providing the robust technical infrastructure and expertise necessary for data hosting and seamless computational workflows.

## AUTHOR CONTRIBUTIONS

D.W.P.T. contributed to the methodology, algorithm development, data analysis, visualization, and manuscript editing. W.K. was involved in conceptualization, algorithm development, data analysis, manuscript editing, and supervision of the work. S.J.A., Z.M.W., N.Z.X.Y., and K.A. contributed to algorithm development. Y.W.L. and E.H. performed experimental validation and investigations. F.T.W., and Y.H.L. supervised the work, conceptualized the study, and edited the manuscript. Y.H.L. wrote the original draft.

## REFERENCES

1 Butler, M. S. J. Nat. Prod. 67, 2141–2153 (2004).

2 Jensen, P. R., Chavarria, K. L., Fenical, W., Moore, B. S. & Ziemert, N. J. Ind. Microbiol. Biotechnol. 41, 203–209 (2014).

3 Poynton, E. F. et al. Nucleic Acids Res. 53, D691–D699 (2024).

4 Lin, Z. et al. Science 379, 1123–1130 (2023).

5 Johnson, S. R. & Lange, B. M. Front. Bioeng. Biotechnol. 3 (2015).

6 Wang, M. et al. Nat. Biotechnol. 34, 828–837 (2016).

7 Tay, D. W. P., Yeo, N. Z. X., Adaikkappan, K., Lim, Y. H. & Ang, S. J. Sci. Data 10, 296 (2023).

8 Goldman, S., Li, J. & Coley, C. W. Anal. Chem. 96, 3419–3428 (2024).

9 Wang, F. et al. Anal. Chem. 93, 11692–11700 (2021).

10 Wei, J. N., Belanger, D., Adams, R. P. & Sculley, D. ACS Cent. Sci. 5, 700–708 (2019).

11 Tay, D. W. P. et al. Sci. Data 11, 977 (2024).

12 Ng, S. B. et al. Nat. Biotechnol. 36, 570–573 (2018).

13 Tay, D. W. P. et al. Commun. Biol. 7, 50 (2024).

14 Chen, J.-Y. et al. Front. Bioeng. Biotechnol. 13 (2025).

15 Liu, W. et al. Nat. Commun. 15, 2775 (2024).

16 Johnson, S. R., Peshwa, M. & Sun, Z. eLife 12, RP91415 (2024).

17 Douze, M. et al. arXiv (2024).

18 Huang, F. et al. Org. Biomol. Chem. 3, 1410–1418 (2005).

19 Bolger, A. M., Lohse, M. & Usadel, B. Bioinformatics 30, 2114–2120 (2014).

20 Hyatt, D. et al. BMC Bioinformatics 11, 119 (2010).

21 Sorokina, M., Merseburger, P., Rajan, K., Yirik, M. A. & Steinbeck, C. J. Cheminform. 13, 2 (2021).

22 Nicolescu, T. O. Mass Spectrometry, 23–78 (2017).

23 Tay, D. W. P., Adaikkappan, K., Yeo, N. Z. X., Ang, S. J. & Lim, Y. H. SG Patent Application No. 10202403057W (2024).

