## Supplementary Information for "Accelerating Natural Product Discovery with Linked MS-Genomics and Language/Transformer-Based Models"

### Workflow for Intelligent Structural Elucidation (WISE)

**Table S1. Technology benchmarking against known existing solutions for structural identification from MS/MS.**

| Tool | **NEIMS[1]** | **CFM-ID 4.0[2]** | **ICEBERG[3]** | **WISE[4,5]** |
|---|---|---|---|---|
| Year | 2019 | 2022 | 2024 | 2024 |
| Creator | Google | Univ. of Alberta | MIT | A*STAR |
| MSMS Similarity to Experimental | 36% | 25% | 42% | 75% |
| MSMS-to-Structure Accuracy* | 8.6% | 8.6% | 12.9% | 42% |
| Novel Structures | X | X | X | ✓ |
| Reference | *ACS Cent. Sci.* **2019**, 5, 700-708 | *Anal. Chem.* **2021**, 23, 11692-11700 | *Anal. Chem.* **2024**, 96, 3419-3428 | SG Patents 10202400652R **2024** and 10202403057W **2024** |

*Top 1 accuracy. Literature (NEIMS, CFM-ID 4.0, ICEBERG) MSMS-to-Structure accuracies taken from Supporting Information of *Anal. Chem.* **2024**, 96, 3419-3428. WISE MSMS-to-Structure accuracy from evaluating 4,505 samples from the public libraries of GNPS Community, NIH round 1, and NIH round 2 taken from GNPS[6].

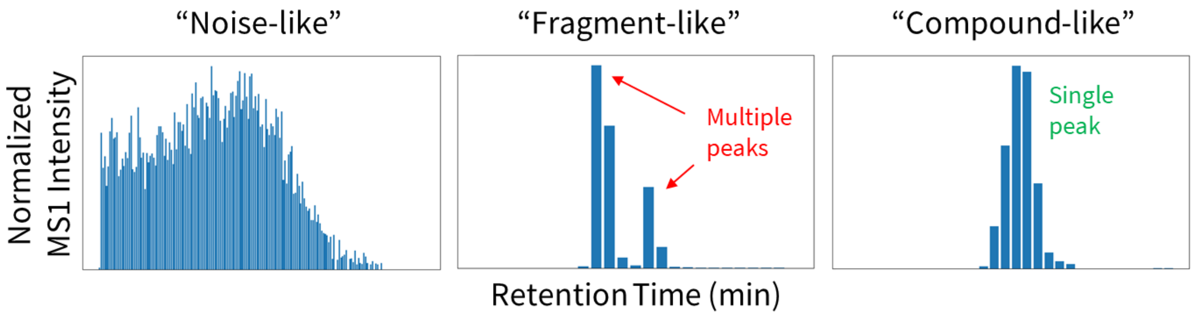

**Figure S1.** Graphs of normalized MS1 intensity of selected m/z’s (mass-to-charge ratio) against retention time (min) for three different signal types – “noise-like”, “fragment-like”, and “compound-like”. Deconvolution analysis done at the individual m/z level, identifying compounds based on their elution pattern over time.

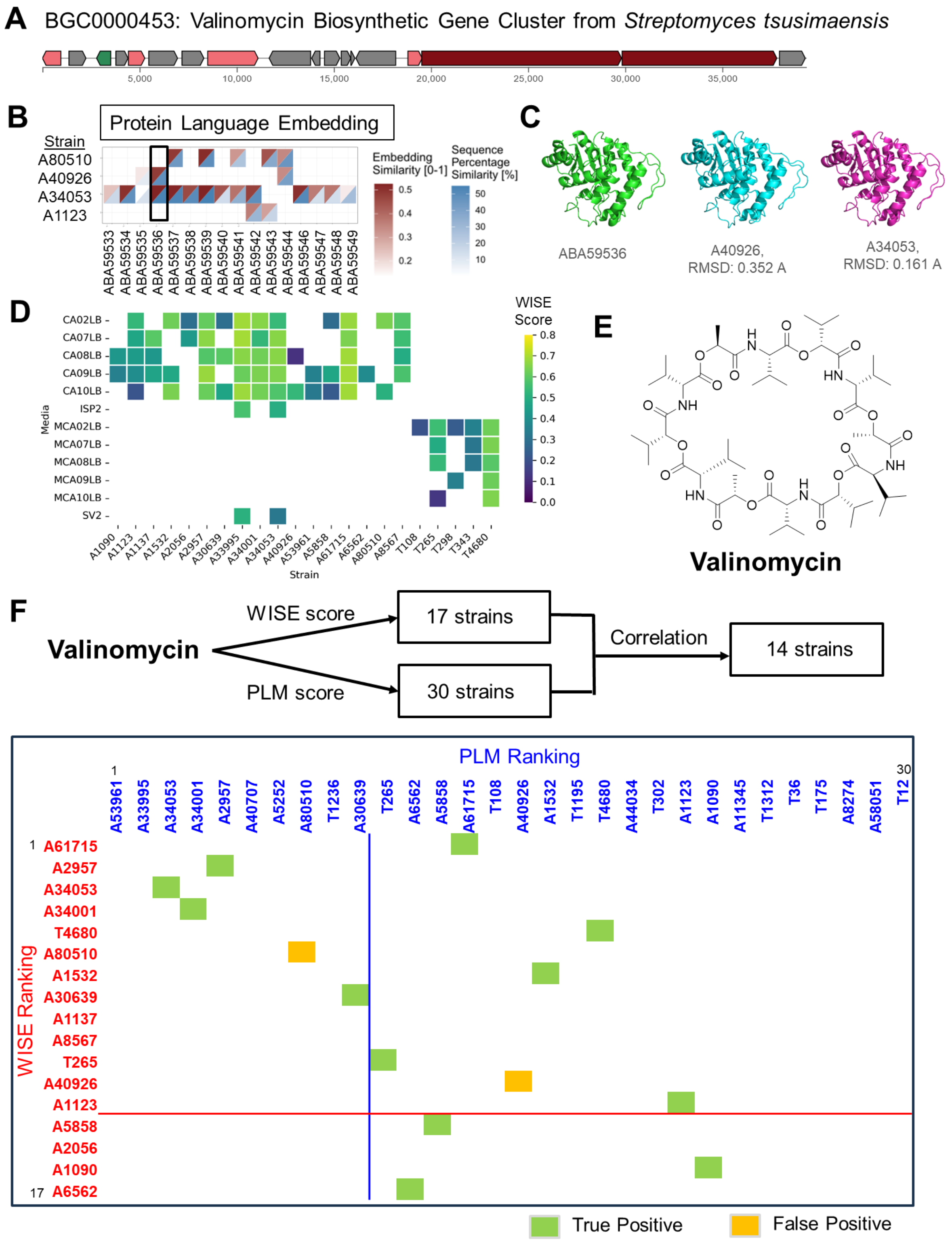

**Figure S2. Case Study: valinomycin. (A)** Elucidated biosynthetic gene cluster BGC0000453, responsible for valinomycin production. **(B)** Similarity (%) of each embedded protein and sequence similarity (%) of the top-scoring protein matches. **(C)** Predicted protein structures of the top protein matches and reductase (ABA59536.1)

from valinomycin BGC. **(D)** Heatmap of WISE scores for strain-media combinations derived from the liquid chromatography-tandem mass spectrometry (LC-MS/MS) data of fermentation extracts. Only strains with at least 1 media combination with non-zero WISE score are shown. Activated strains (if any) are referred to by their parent strain instead of individually (details of the activated strains are in **Table S6**) **(E)** Structure of valinomycin. **(F)** Workflow of the co-occurrence strategy and matrix ranking table with 35% WISE score and 0.75 PLM score cutoff. Strains that have been cross-validated are highlighted in green or orange, indicating whether these compounds were present (green) or absent (orange). Lines indicating higher confidence thresholds of 50% WISE score (in red) and 2.5 PLM score (in blue) are also included.

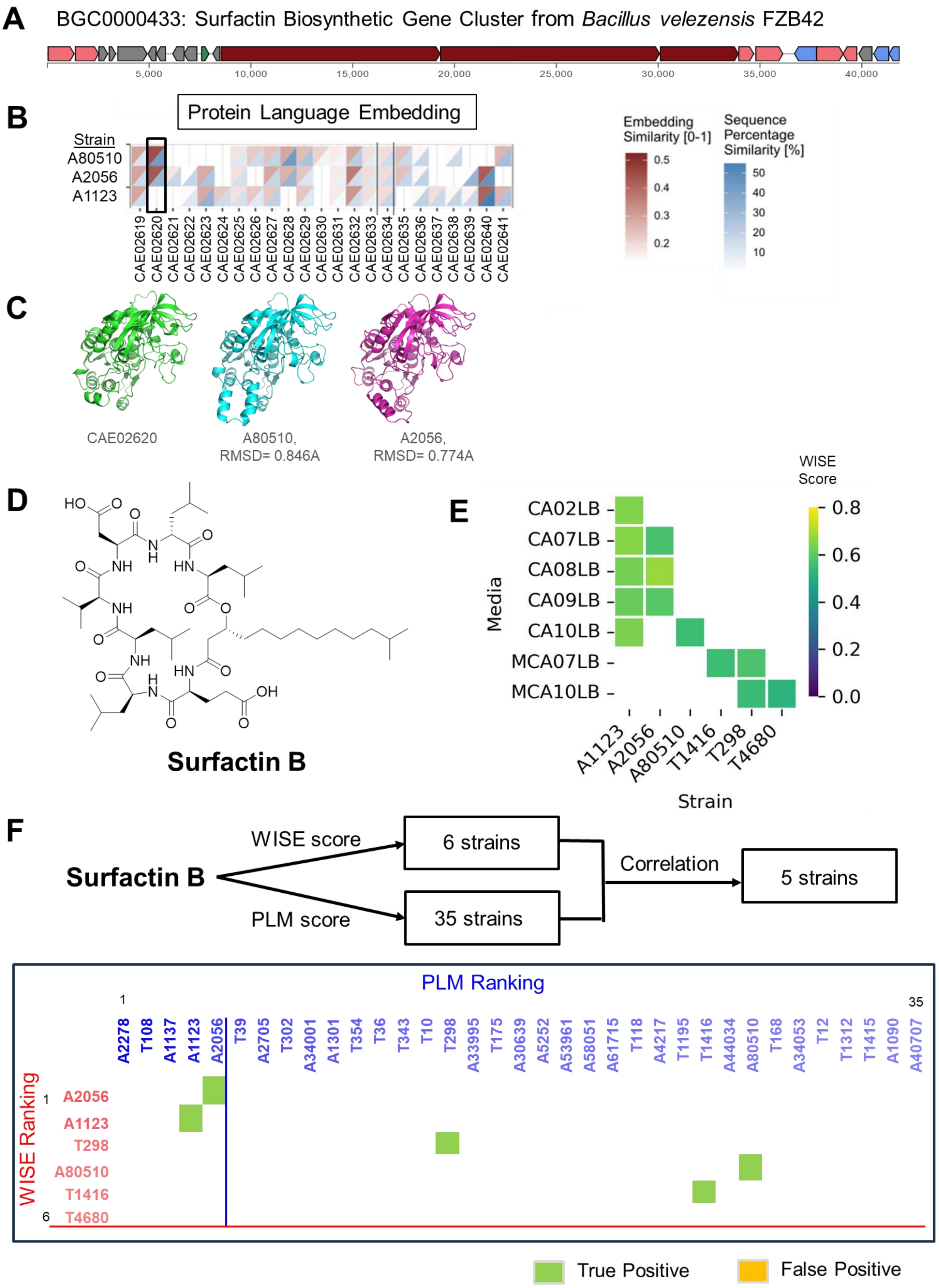

**Figure S3. Case Study: surfactin B. (A)** Elucidated biosynthetic gene cluster BGC0000433, responsible for surfactin production. **(B)** Similarity (%) of each embedded protein and sequence similarity (%) of the top-scoring protein matches. **(C)** Predicted protein structures of the top protein matches and alcohol dehydrogenase (CAE02620.1) from surfactin BGC. **(D)** Structure of surfactin B. **(E)** Heatmap of WISE scores for strain-media combinations derived from the liquid chromatography-tandem mass spectrometry (LC-MS/MS) data of fermentation extracts. Only strains with at least 1 media combination with non-zero WISE score are shown. Activated strains (if any) are referred to by their parent strain instead of individually (details of the activated strains are in **Table S7**). **(F)** Workflow of the co-occurrence strategy and matrix ranking table with 35% WISE score and 0.75 PLM score cut-off. Strains that have been cross-validated are highlighted in green or orange, indicating whether these compounds were present (green) or absent (orange). Lines indicating higher confidence thresholds of 50% WISE score (in red) and 2.5 PLM score (in blue) are also included.

#### Genomic Data

**Table S2. Representative Statistics on Genomic dataset.**

| Strain | Number of scaffolds | Total length (b) | GC (%) | n50 | l50 | Number of Proteins |
|---|---|---|---|---|---|---|
| T1236 | 30012 | 15357158 | 68.0 | 720 | 2246 | 35457 |
| A44034 | 7092 | 8626198 | 71.0 | 4037 | 534 | 13256 |
| A1123 | 11815 | 10444413 | 65.5 | 2343 | 1096 | 17850 |
| A40926 | 35583 | 19007065 | 68.5 | 684 | 1755 | 43613 |
| A34053 | 35890 | 19126044 | 68.5 | 656 | 2066 | 42948 |
| T265 | 35500 | 17328132 | 68.3 | 525 | 3405 | 41217 |
| A58051 | 55310 | 26899034 | 69.1 | 458 | 6808 | 65217 |
| A2056 | 28852 | 21229529 | 68.9 | 4380 | 887 | 41021 |
| T354 | 26504 | 15028253 | 68.3 | 720 | 2439 | 32476 |
| A80510 | 62516 | 29872159 | 68.5 | 467 | 7510 | 73396 |
| T343 | 19735 | 13060844 | 68.6 | 2058 | 1063 | 25905 |

#### WISE Processed Results for Case Studies

**Table S3.** Global regulators

| Regulator | Description | Accession code |
|---|---|---|
| AdpA | AraC/XylS transcriptor family protein | SCO 2792 |
| FAS | Fatty acyl CoA synthase | SCO 6196 |
| Crp | Cyclic AMP receptor protein | SCO 3571 |
| SarA | Putative membrane protein gene | SCO 4069 |
| RedD | AfsR/SARP family transcriptor regulator, RedD | SLIV 09220 |

#### MS/MS Spectral Comparisons Between Top Consensus Matches and GNPS References for Case Studies

**Table S4. Metadata on Top Consensus Matches and GNPS References**

| Compound | Strain ID | Parent | Media | WISE Score | GNPS Ref |
|---|---|---|---|---|---|
| Valinomycin | A2957 | A2957 | CA09LB | 68% | CCMSLIB00005721598 |
| Valinomycin | A34053 | A34053 | CA07LB | 66% | CCMSLIB00005721598 |
| Valinomycin | A34001 | A34001 | CA02LB | 65% | CCMSLIB00005721598 |
| Surfactin B | A2056 | A2056 | CA08LB | 70% | CCMSLIB00005727861 |
| Surfactin B | A100214 | A1123 | CA07LB | 67% | CCMSLIB00005727861 |
| Neomycin B | T354 | T354 | MCA09LB | 44% | CCMSLIB00010114511 |
| Neomycin B | T10063 | T1236 | MCA08LB | 40% | CCMSLIB00010114511 |
| Neomycin B | T10090 | T343 | MCA02LB | 39% | CCMSLIB00010114511 |

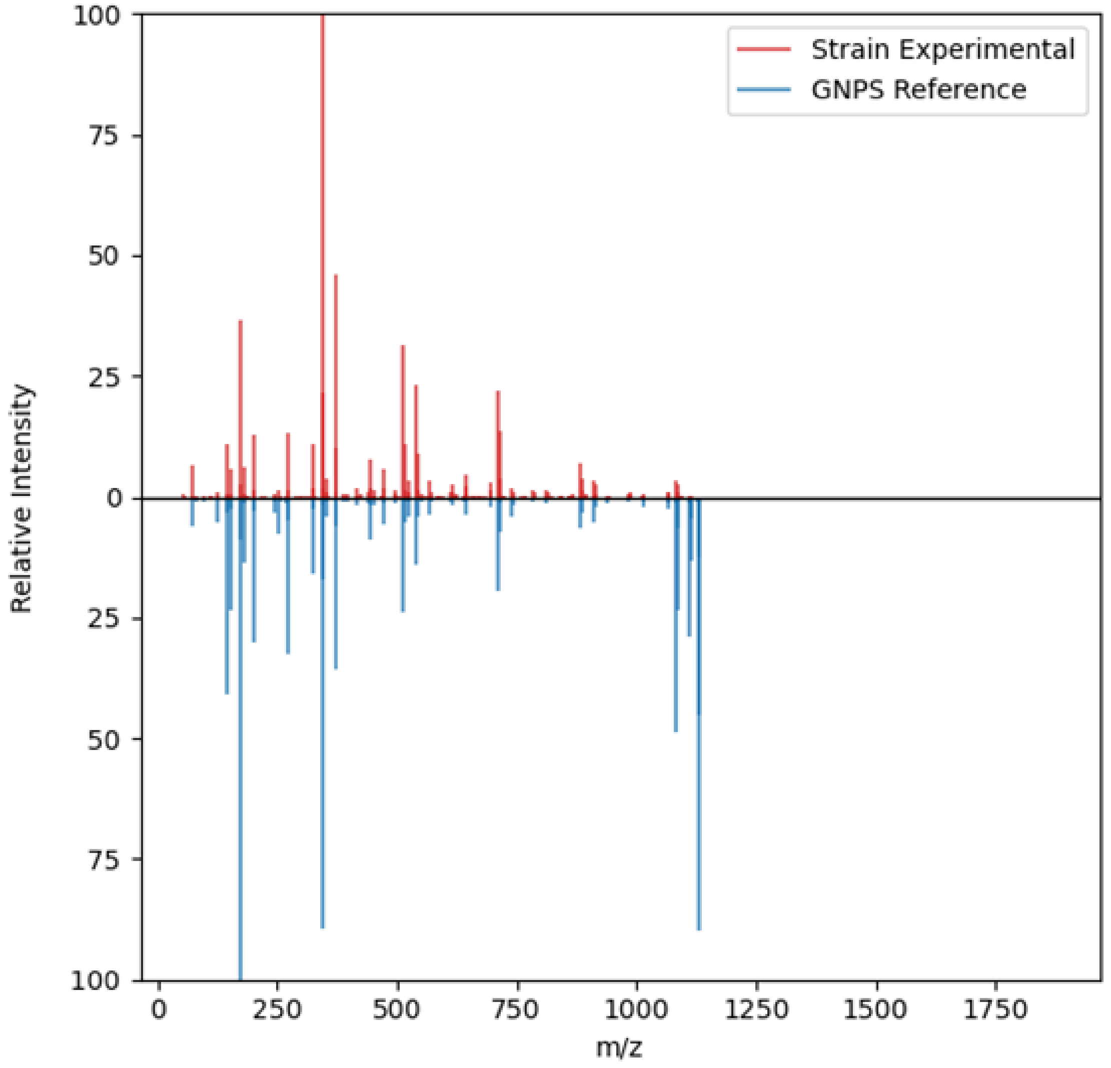

**Figure S7.** MS/MS spectral comparison of valinomycin found from fermentation of strain A2957 in CA09LB media versus GNPS reference.

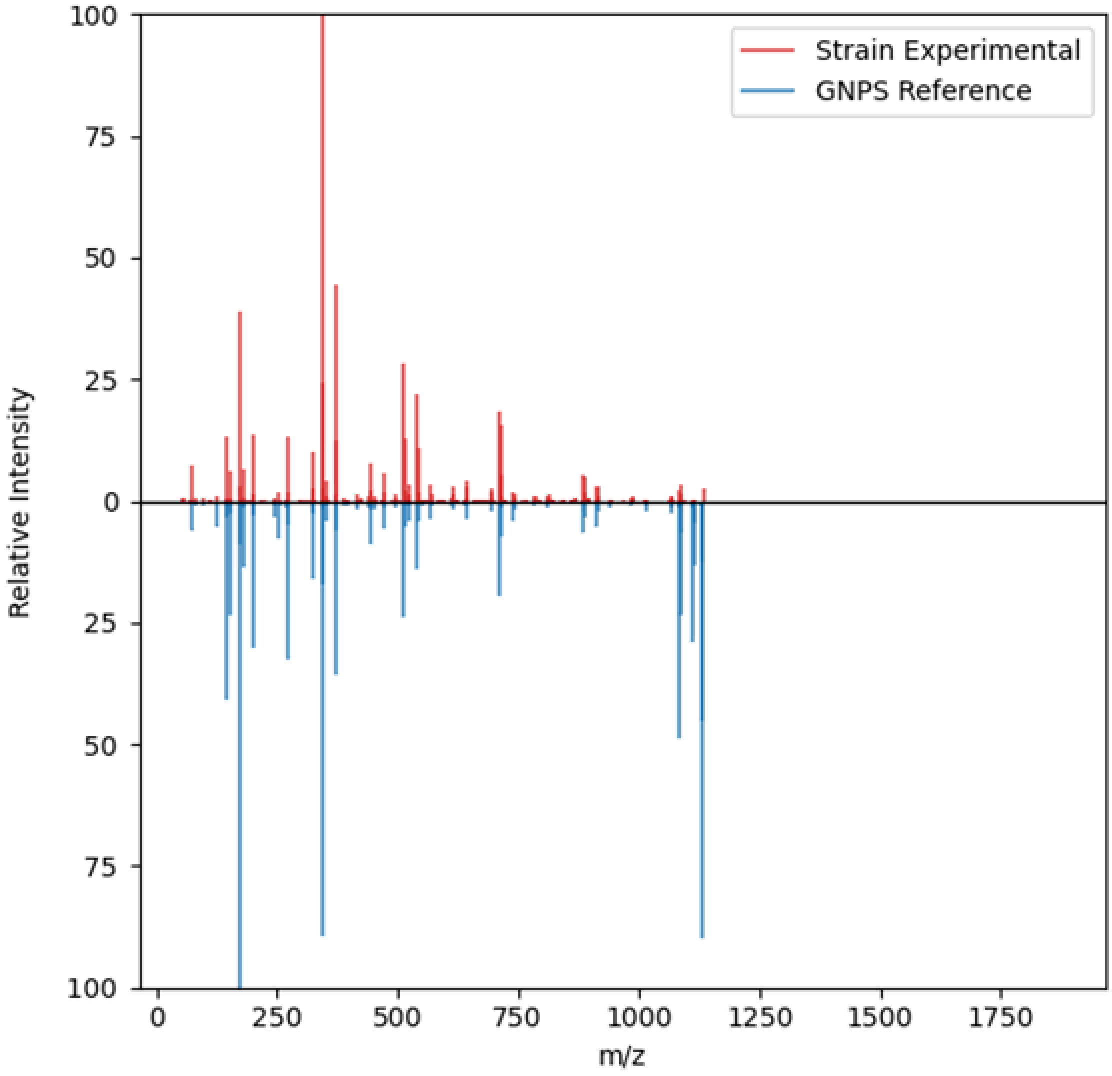

**Figure S8.** MS/MS spectral comparison of valinomycin found from fermentation of strain A34053 in CA07LB media versus GNPS reference.

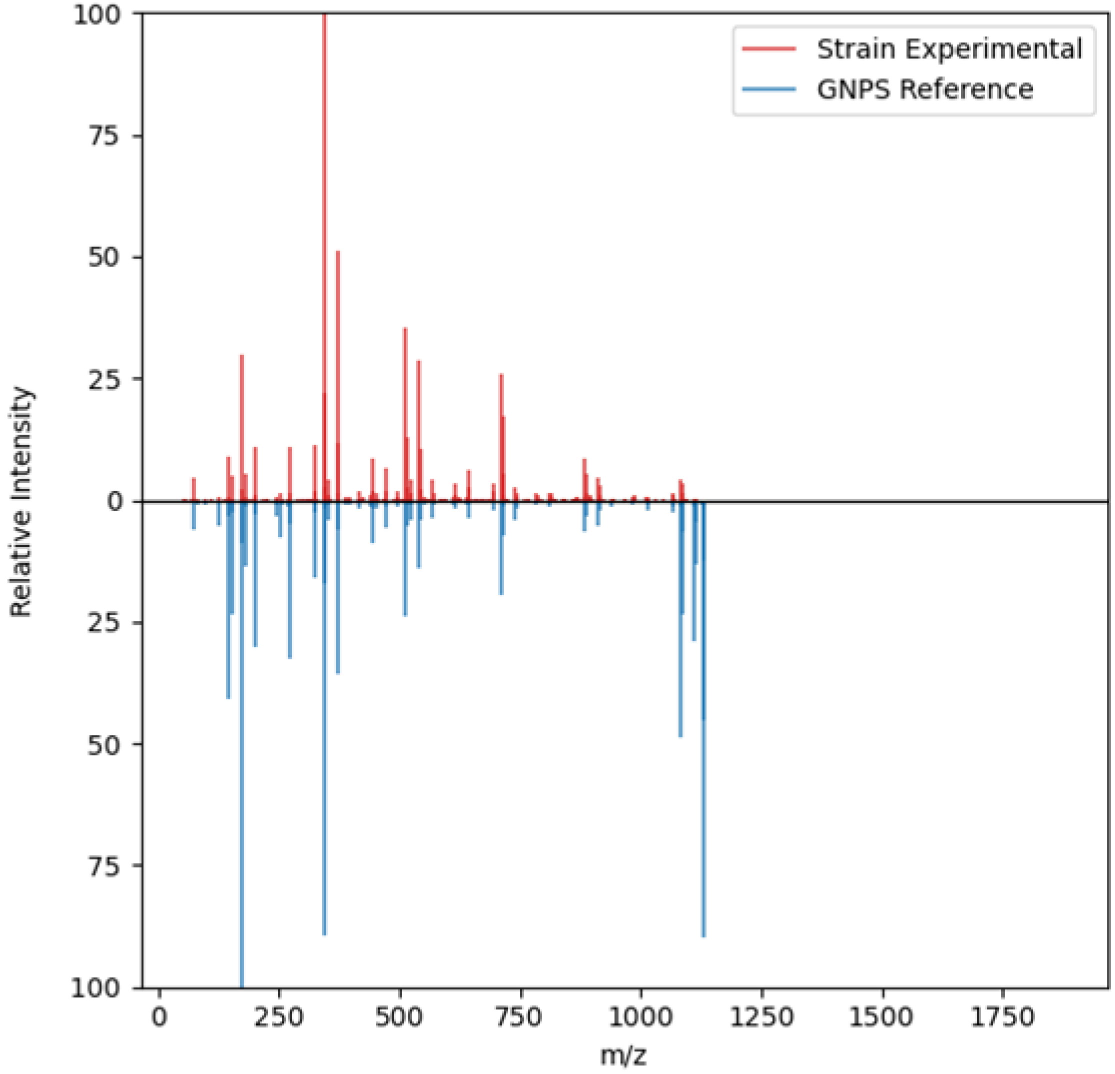

**Figure S9.** MS/MS spectral comparison of valinomycin found from fermentation of strain A34001 in CA02LB media versus GNPS reference.

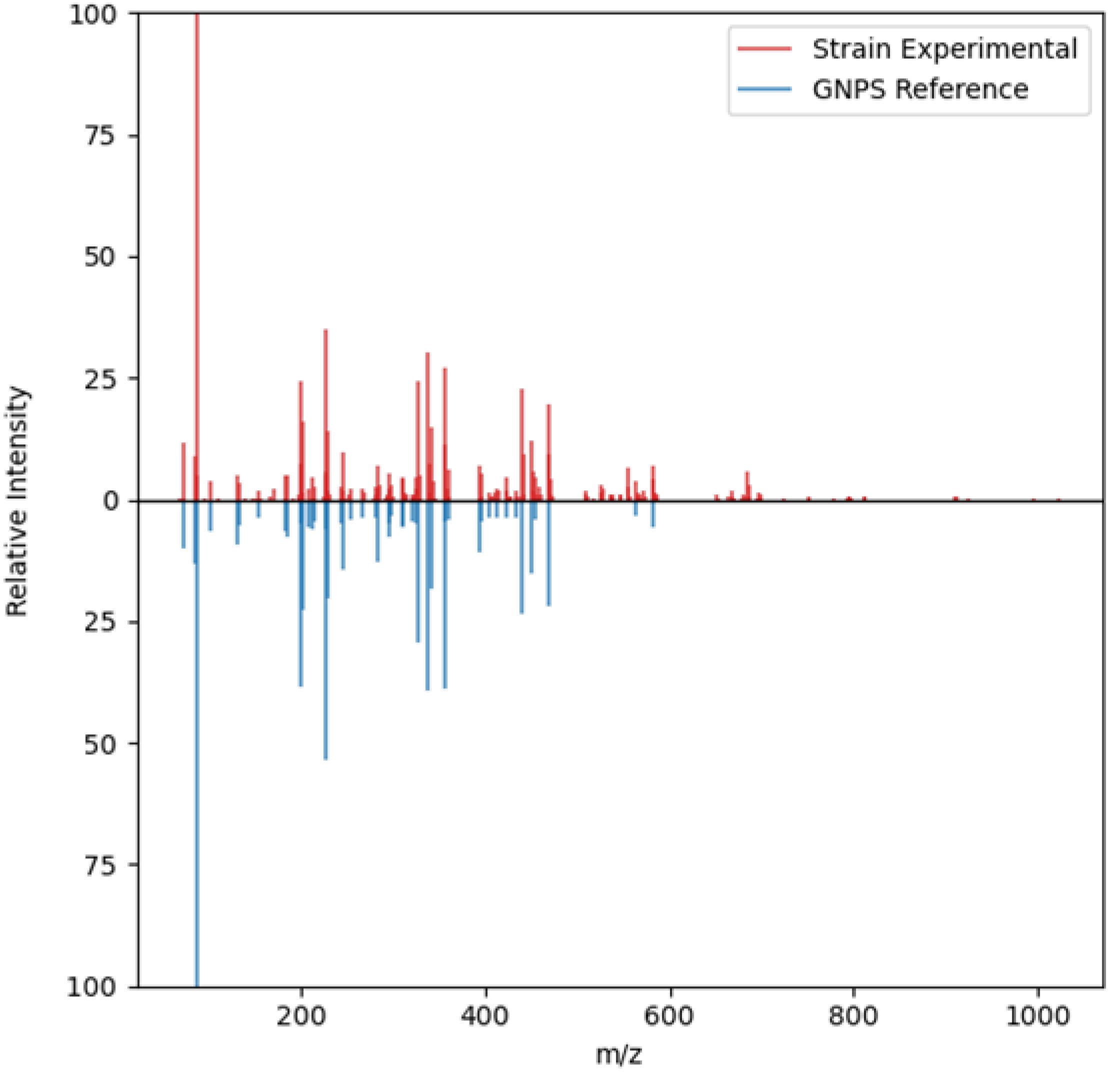

**Figure S10.** MS/MS spectral comparison of surfactin B found from fermentation of strain A2056 in CA08LB media versus GNPS reference.

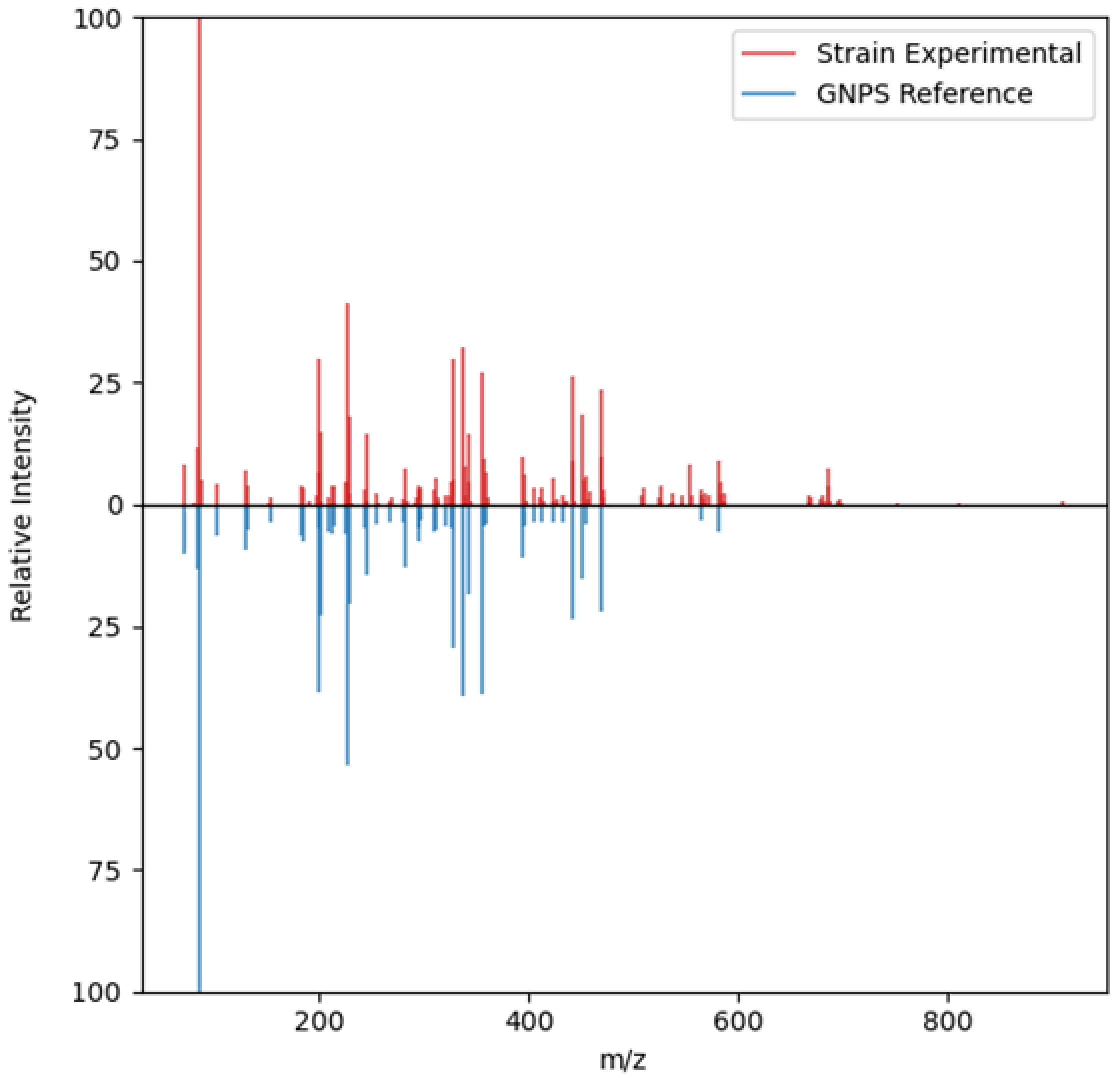

**Figure S11.** MS/MS spectral comparison of surfactin B found from fermentation of strain A100214 in CA07LB media versus GNPS reference.

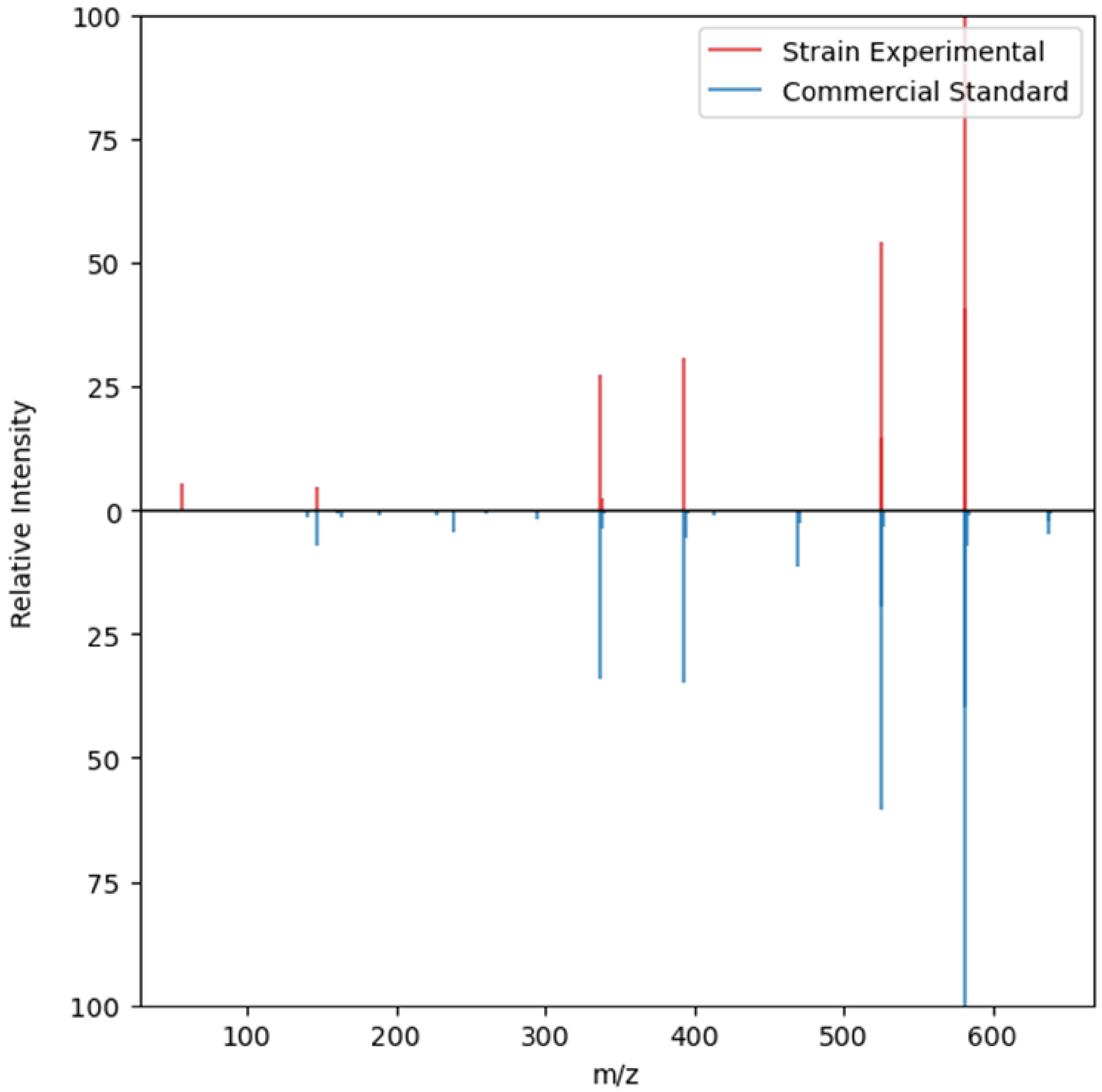

**Figure S12.** MS/MS spectral comparison of neomycin B found from fermentation of strain T354 in MCA09LB media versus commercial reference.

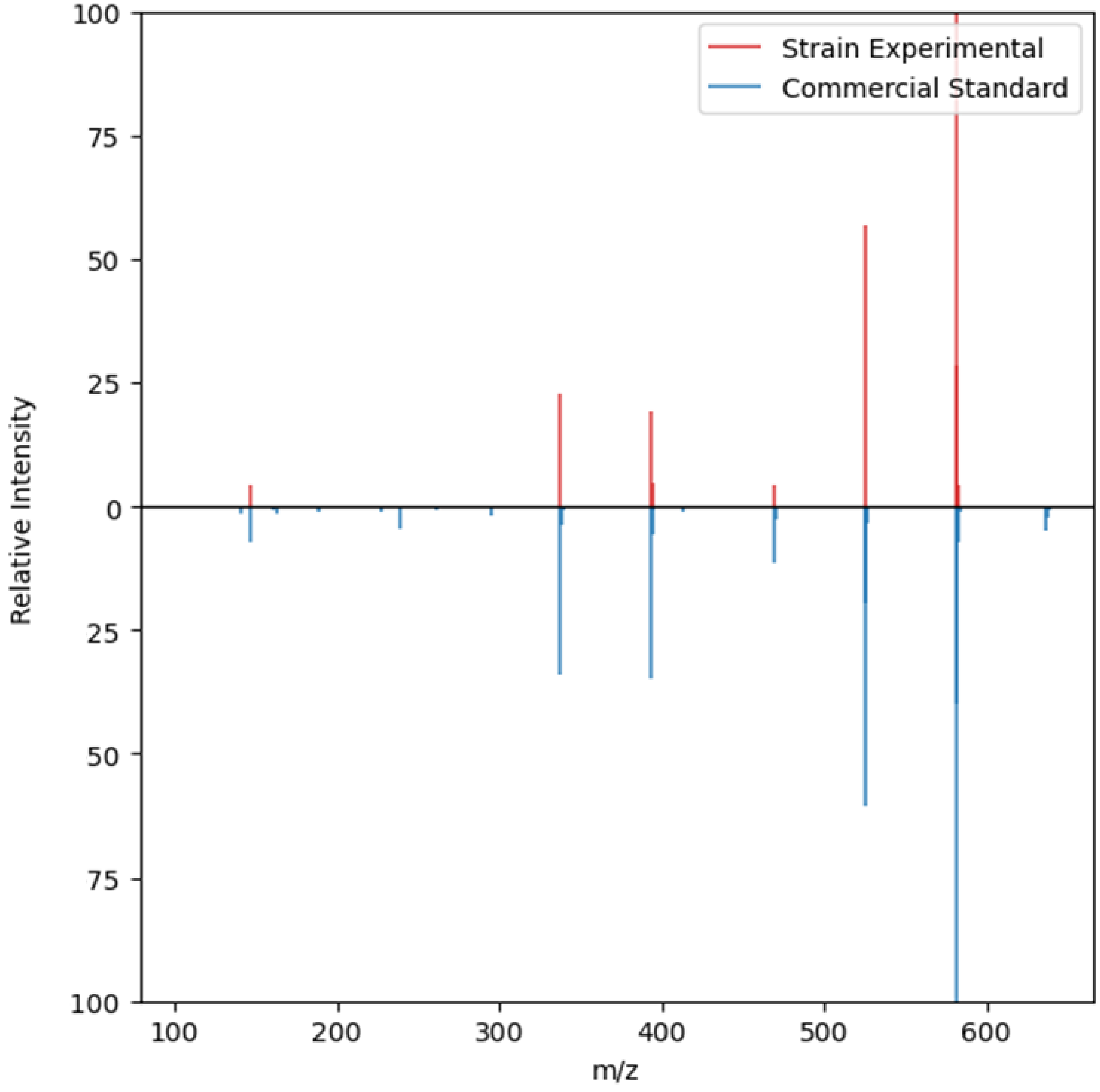

**Figure S13.** MS/MS spectral comparison of neomycin B found from fermentation of strain T10063 in MCA08LB media versus commercial reference.

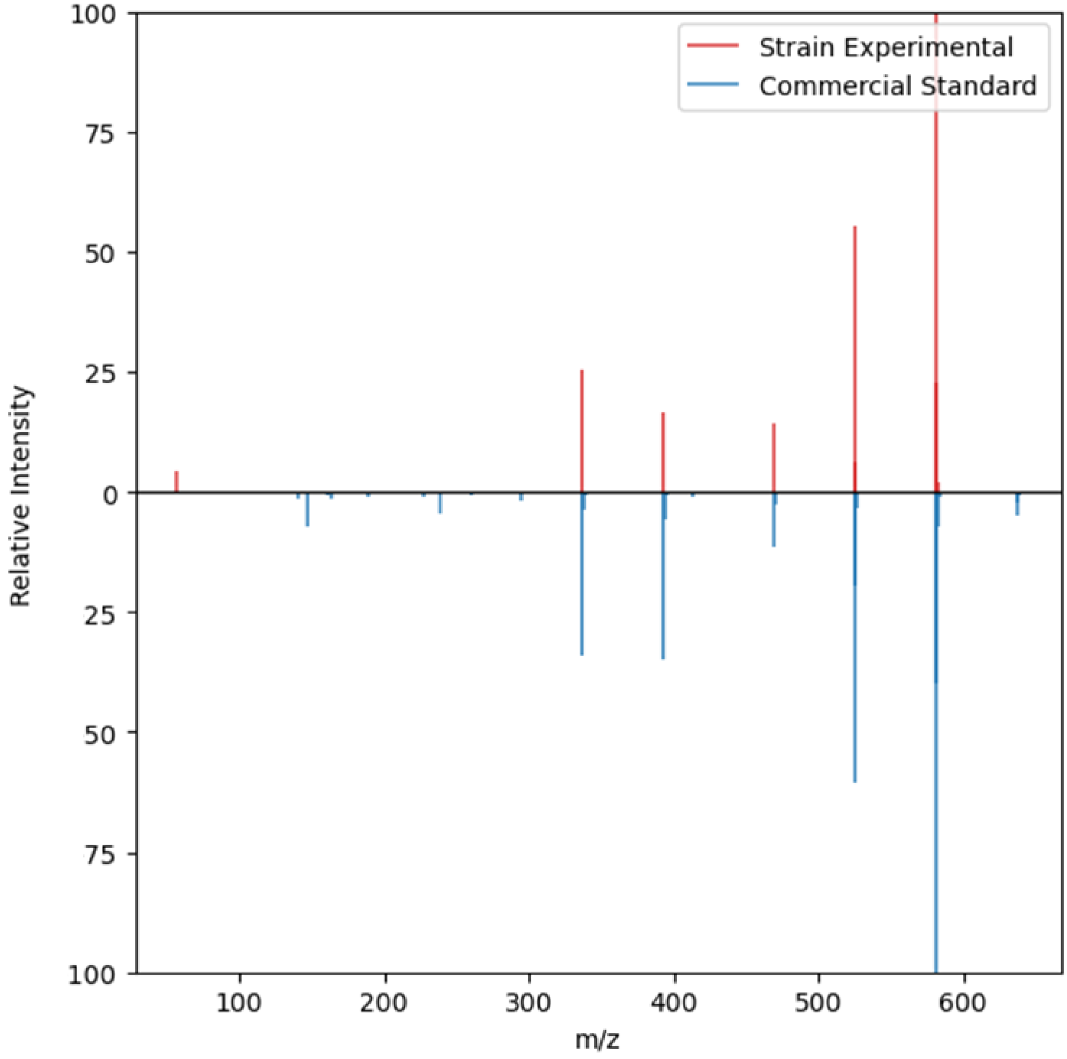

**Figure S14.** MS/MS spectral comparison of neomycin B found from fermentation of strain T10090 in MCA02LB media versus commercial reference.

#### Strain Information

**Table S5.** Strain ID, its associated taxonomy and references of previous observations

| **Strain ID** | **Identity % based on rRNA BLAST** | **References** |
|---|---|---|
| Surfactin producers | | |
| A2056 | *Streptomyces ardesiacus*, 98.2% | This study |
| A1123 | *Thermoactinomyces vulgaris*, 100% | This study |
| T298 | *Streptomyces fradiae*, 99.69% | This study |
| A80510 | *Micromonospora maritima*, 97.18% | [7] |
| T1416 | *Streptomyces tendae*, 99.21% | [8] |
| Neomycin producers | | |
| T354 | *Streptomyces sampsonii,* 99.93% | This study |
| T1236 | *Streptomyces ardesiacus*, 99.91% | This study |
| T343 | *Streptomyces albogriseolus*, 100% | https://en.wikipedia.org/wiki/Streptomyces_albogriseolus, accessed 9 Jan 2025 |

#### Abundance of Case Study Compounds

**Table S6.** Comparison of MS abundance among selected Valinomycin producers from the 12 parent strains found via co-occurrence of WISE and PLM scores.

| Parent strain | Strain ID | Media | Regulator | WISE Score | MS Abundance |
|---|---|---|---|---|---|
| A1090 | A1090 | - | - | N.F. | - |
| A1090 | A100190 | CA08LB | Crp | 42% | 6.83E+05 |
| A1123 | A1123 | CA02LB | - | 52% | 1.33E+06 |
| A1123 | A100131 | CA08LB | SarA | 44% | 3.26E+06 |
| A1532 | A1532 | - | - | N.F. | - |
| A1532 | A100281 | CA10LB | FAS | 64% | 1.89E+07 |
| A1532 | A100281 | CA02LB | FAS | 62% | 2.15E+07 |
| A2957 | A2957 | CA09LB | - | 68% | 1.54E+05 |
| A2957 | A100261 | CA09LB | RedD | 63% | 1.83E+07 |
| A30639 | A30639 | CA08LB | - | 59% | 3.36E+05 |
| A30639 | A100312 | CA10LB | Crp | 47% | 4.72E+05 |
| A34001 | A34001 | CA02LB | - | 65% | 4.58E+05 |
| A34001 | A100267 | CA09LB | Crp | 48% | 3.80E+07 |
| A34053 | A34053 | CA07LB | - | 66% | 2.04E+05 |
| A34053 | A100034 | CA09LB | Crp | 64% | 2.47E+07 |
| A5858 | A5858 | CA09LB | - | 47% | 3.84E+05 |
| A61715 | A61715 | CA07LB | - | 52% | 8.33E+06 |
| A61715 | A100153 | CA09LB | Crp | 69% | 2.00E+07 |
| A61715 | A100157 | CA07LB | FAS | 51% | 2.86E+07 |
| A6562 | A6562 | CA09LB | - | 38% | 2.97E+05 |
| T265 | T265 | - | - | N.F. | - |
| T265 | T10034 | MCA02LB | Crp | 55% | 1.51E+05 |
| T265 | T10051 | MCA08LB | RedD | 50% | 1.68E+06 |
| T4680 | T4680 | MCA10LB | - | 41% | 6.04E+05 |
| T4680 | T10016 | MCA10LB | Crp | 64% | 3.14E+06 |
| T4680 | T10018 | MCA10LB | Crp | 55% | 4.17E+07 |

N.F. = not found

**Table S7**. Comparison of MS abundance among selected Surfactin B producers from the 5 parent strains found via co-occurrence of WISE and PLM scores.

| Parent strain | Strain ID | Media | Regulator | WISE Score | MS Abundance |
|---|---|---|---|---|---|
| A1123 | A1123 | - | - | N.F. | - |
| A1123 | A100214 | CA07LB | RedD | 67% | 2.08E+06 |
| A1123 | A100214 | CA08LB | RedD | 47% | 3.45E+06 |
| A2056 | A2056 | CA08LB | - | 70% | 3.72E+06 |
| A80510 | A80510 | CA10LB | - | 57% | 3.58E+05 |
| T1416 | T1416 | - | - | N.F. | - |
| T1416 | T10217 | MCA07LB | FAS | 57% | 1.30E+05 |
| T1416 | T10225 | MCA07LB | RedD | 56% | 5.52E+05 |
| T298 | T298 | MCA10LB | - | 52% | 1.19E+05 |
| T298 | T10028 | MCA07LB | RedD | 59% | 4.10E+05 |

N.F. = not found

**Table S8.** Comparison of MS abundance among selected Neomycin B producers from the 3 parent strains found via co-occurrence of WISE and PLM scores.

| Parent strain | Strain ID | Media | Regulator | WISE Score | MS Abundance |
|---|---|---|---|---|---|
| T1236 | T1236 | - | - | N.F. | - |
| T1236 | T10063 | MCA08LB | RedD | 40% | 7.05E+04 |
| T343 | T343 | MCA09LB | - | 35% | 9.23E+04 |
| T343 | T10090 | MCA02LB | RedD | 39% | 1.44E+05 |
| T354 | T354 | MCA09LB | None | 44% | 1.23E+05 |

N.F. = not found

#### References

1. Wei, J. N.; Belanger, D.; Adams, R. P.; Sculley, D., Rapid Prediction of Electron–Ionization Mass Spectrometry Using Neural Networks. *ACS Cent. Sci.* **2019,** *5* (4), 700-708.
2. Wang, F.; Liigand, J.; Tian, S.; Arndt, D.; Greiner, R.; Wishart, D. S., CFM-ID 4.0: More Accurate ESI-MS/MS Spectral Prediction and Compound Identification. *Anal. Chem.* **2021,** *93* (34), 11692-11700.
3. Goldman, S.; Li, J.; Coley, C. W., Generating Molecular Fragmentation Graphs with Autoregressive Neural Networks. *Anal. Chem.* **2024,** *96* (8), 3419-3428.
4. Tay, D. W. P.; Ang, S. J.; Lim, Y. H.; Wong, Z. M. Generation of Collision Energy Specific Tandem Mass Spectra from Molecular Structures. SG Patent Application No. 10202400652R, 2024.
5. Tay, D. W. P.; Adaikkappan, K.; Yeo, N. Z. X.; Ang, S. J.; Lim, Y. H. Automated Workflow for Intelligent Structural Elucidation (WISE) from Raw Liquid Chromatography Tandem Mass Spectrometry Data. SG Patent Application No. 10202403057W, 2024.
6. Wang, M.; Carver, J. J.; Phelan, V. V.; Sanchez, L. M.; Garg, N.; Peng, Y.; Nguyen, D. D.; Watrous, J.; Kapono, C. A.; Luzzatto-Knaan, T.; Porto, C.; Bouslimani, A.; Melnik, A. V.; Meehan, M. J.; Liu, W.-T.; Crüsemann, M.; Boudreau, P. D.; Esquenazi, E.; Sandoval-Calderón, M.; Kersten, R. D.; Pace, L. A.; Quinn, R. A.; Duncan, K. R.; Hsu, C.-C.; Floros, D. J.; Gavilan, R. G.; Kleigrewe, K.; Northen, T.; Dutton, R. J.; Parrot, D.; Carlson, E. E.; Aigle, B.; Michelsen, C. F.; Jelsbak, L.; Sohlenkamp, C.; Pevzner, P.; Edlund, A.; McLean, J.; Piel, J.; Murphy, B. T.; Gerwick, L.; Liaw, C.-C.; Yang, Y.-L.; Humpf, H.-U.; Maansson, M.; Keyzers, R. A.; Sims, A. C.; Johnson, A. R.; Sidebottom, A. M.; Sedio, B. E.; Klitgaard, A.; Larson, C. B.; Boya P, C. A.; Torres-Mendoza, D.; Gonzalez, D. J.; Silva, D. B.; Marques, L. M.; Demarque, D. P.; Pociute, E.; O'Neill, E. C.; Briand, E.; Helfrich, E. J. N.; Granatosky, E. A.; Glukhov, E.; Ryffel, F.; Houson, H.; Mohimani, H.; Kharbush, J. J.; Zeng, Y.; Vorholt, J. A.; Kurita, K. L.; Charusanti, P.; McPhail, K. L.; Nielsen, K. F.; Vuong, L.; Elfeki, M.; Traxler, M. F.; Engene, N.; Koyama, N.; Vining, O. B.; Baric, R.; Silva, R. R.; Mascuch, S. J.; Tomasi, S.; Jenkins, S.; Macherla, V.; Hoffman, T.; Agarwal, V.; Williams, P. G.; Dai, J.; Neupane, R.; Gurr, J.; Rodríguez, A. M. C.; Lamsa, A.; Zhang, C.; Dorrestein, K.; Duggan, B. M.; Almaliti, J.; Allard, P.-M.; Phapale, P.; Nothias, L.-F.; Alexandrov, T.; Litaudon, M.; Wolfender, J.-L.; Kyle, J. E.; Metz, T. O.; Peryea, T.; Nguyen, D.-T.; VanLeer, D.; Shinn, P.; Jadhav, A.; Müller, R.; Waters, K. M.; Shi, W.; Liu, X.; Zhang, L.; Knight, R.; Jensen, P. R.; Palsson, B. Ø.; Pogliano, K.; Linington, R. G.; Gutiérrez, M.; Lopes, N. P.; Gerwick, W. H.; Moore, B. S.; Dorrestein, P. C.; Bandeira, N., Sharing and community curation of mass spectrometry data with Global Natural Products Social Molecular Networking. *Nat. Biotechnol.* **2016,** *34* (8), 828-837.
7. Pang, X.; Zhao, J.; Fang, X.; Liu, H.; Zhang, Y.; Cen, S.; Yu, L., Surfactin derivatives from Micromonospora sp. CPCC 202787 and their anti-HIV activities. *J. Antibiot.* **2017,** *70* (1), 105-108.
8. Richter, M.; Willey, J. M.; Süßmuth, R.; Jung, G.; Fiedler, H.-P., Streptofactin, a novel biosurfactant with aerial mycelium inducing activity from Streptomyces tendae Tü 901/8c. *FEMS Microbiol. Lett.* **1998,** *163* (2), 165-171.
